# Probing brain developmental patterns of myelination and associations with psychopathology in youth using gray/white matter contrast

**DOI:** 10.1101/305995

**Authors:** Linn B. Norbom, Nhat Trung Doan, Dag Alnæs, Tobias Kaufmann, Torgeir Moberget, Jaroslav Rokicki, Ole A. Andreassen, Lars T. Westlye, Christian K. Tamnes

## Abstract

**Background:** Cerebral myeloarchitecture shows substantial development across childhood and adolescence, and aberrations in these trajectories are relevant for a range of mental disorders. Differential myelination between intracortical and subjacent white matter can be approximated using signal intensities in T1-weighted magnetic resonance images (MRI).

**Methods:** To test the sensitivity of gray/white matter contrast (GWC) to age and individual differences in psychopathology and general cognitive ability in youth (8-23 years), we formed data-driven psychopathology and cognitive components using a large population-based sample, the Philadelphia Neurodevelopmental Cohort (PNC) (n=6487, 52% females). We then tested for associations with regional GWC defined by an independent component analysis (ICA) in a subsample with available MRI data (n=1467, 53% females).

**Results:** The analyses revealed a global GWC component, which showed an age-related decrease from late childhood and across adolescence. In addition, we found regional anatomically meaningful components with differential age associations explaining variance beyond the global component. When accounting for age and sex, both higher symptom levels of anxiety or prodromal psychosis and lower cognitive ability were associated with higher GWC in insula and cingulate cortices and with lower GWC in pre- and postcentral cortices. We also found several additional regional associations with anxiety, prodromal psychosis and cognitive ability.

**Conclusion:** Independent modes of GWC variation are sensitive to global and regional brain developmental processes, possibly related to differences between intracortical and subjacent white matter myelination, and individual differences in regional GWC are associated with both mental health and general cognitive functioning.

## Introduction

Adolescence is a time of extensive changes in the sociocultural domain, as well as within body, cognition, emotion and behavior (1, 2). Brain development during this period involves multiple biological processes that dynamically interact with the environment, and shows temporal and spatial heterogeneity across tissue types, measures and individuals (3-6). Increased knowledge about these brain developmental patterns and their individual differences is key to inform ontogenetic models of psychopathology.

Although current psychiatric diagnostic tools rely on manifest symptoms that often appear relatively late in illness progression, numerous mental disorders are considered to have a neurodevelopmental origin (7, 8), with additional risk factors related to social behavior and adverse effects of drug and alcohol use (9). Moreover, common symptoms of mental illness form spectra in the general population (10-13). Neuroimaging studies using population based youth samples may thus provide early identification of individuals at risk and insight into the development of mental disorders, and discern brain phenotypes associated with the full range of variation in symptoms of mental illness.

The information processing analogy of the brain has provided a useful framework for the study of brain aberrations in clinical populations (14). An underlying assumption is that a healthy brain is characterized by efficient communication between regions, made possible by a structural backbone of myelinated axons and pathways. Myelin is key for efficient neural signaling as it increases speed and reliability of the nerve signal, provides support and prevents aberrant sprouting of nerve connections (15, 16). Neuroimaging studies have described developmental trajectories (3, 17) and case-control differences (18) in the organization and microstructure of these white matter (WM) pathways. A recent finding in an overlapping sample, indicate that frontotemporal WM dysconnectivity is a transdiagnostic brain phenotype, associated with both higher levels of psychopathology and lower cognitive abilities (19).

Importantly, beyond WM, the cerebral cortex is also myelinated. Intracortical myelin is predominantly found in deeper layers of cortex, in part due to proliferation of myelin from WM, penetrating the inner periphery of cortical neuropil (20-22). Intracortical myelination is a crucial feature of brain development (23), of which maturational timing conforms with a general posterior-anterior gradient, with additional regional specificity as highly myelinated regions, such as sensorimotor and early association cortices, mature early (23, 24). Cortical myelin content, as indirectly assessed *in vivo* using magnetic resonance imaging (MRI), has been linked to cognitive performance in youth (25) and adults (26), and abnormal intracortical myelination is a candidate mechanism for common mental disorders (8, 16, 27, 28). Compared to closely related non-human primates, postnatal intracortical myelination in humans is exceptionally extended (29). Although there are clear benefits to allow for prolonged environmental influences on brain circuit establishment and refinement, this human-specific shift in timing and extension of cortical development may come at a cost, as diverse forms of psychopathology typically emerge during adolescence (30).

Cholesterol in myelin is a major determinant of the intensity in the T1-weighted MRI signal (31, 32), and cortical gray matter (GM) intensity has been shown to correspond closely with histologically based myelin profiles (33), and specifically with myelin rather than iron content (34). The differential myelination of the cerebral cortex and subjacent WM can be approximated using the gray/white matter contrast (GWC) of the T1-signal intensity. Lower GWC indicates that GM and WM intensities are more similar, while a higher contrast reflects a larger discrepancy, however, the biological interpretation is likely complex. GWC exhibits significant heritability (35), and shows regional aging related patterns (36). Although dysmyelination is a viable candidate mechanism for brain network dysfunction, few studies have examined this measure in relation to mental health, and only in adults with psychosis (37-39).

The present study aimed to test the sensitivity of GWC to individual differences in age and psychopathology in youth (8-23 years). We used the Philadelphia Neurodevelopmental Cohort (PNC, n=6487) to develop data-driven clinical and cognitive components (19). In a subsample with available MRI data (n=1467), we performed an independent component analysis (ICA) of vertex wise GWC across the brain surface and tested for associations with age, different psychopathology components, and general cognitive ability. We hypothesized that GWC generally would show a negative association with age, possibly due to protracted myelination of cortex compared to subjacent WM, but also would show regional age-related patterns. Next, we hypothesized that youth with increased symptoms of psychopathology or lower cognitive ability would show regionally higher GWC, possibly indicating lower levels of intracortical myelin compared to subjacent WM.

## Methods and Materials

### Participants

The analyses were based on the publicly available PNC (permission #8642), a large population based sample comprising MRI, cognitive, clinical and genetic data (40, 41). Further details on the project and procedures are described in the Supplemental Materials and elsewhere (40, 42, 43).

As reported previously (19), all individuals (n=6487, 3379 females) made available aged 8-22 years from the PNC, were included for psychopathology and cognitive analyses. Missing clinical item scores were replaced with the nearest neighbor value based on Euclidean distance. We then formed data-driven cognitive and clinical components, and tested for associations with GWC in a subsample of individuals with available and quality controlled MRI data. From 1601 available scans, we excluded participants with severe medical health conditions (n=70), based on a severity index rating by trained personnel in the PNC study team (44), or due to incomplete or poor quality MRI data (n=64, see below). The final MRI sample consisted of 1467 individuals (776 females) aged 8.2-23.2 years. 66 and 65 participants had missing clinical and cognitive data, respectively, and were excluded from relevant analyses. We refer to Supplemental Table S1 for MRI- and psychopathology/cognitive sample demographics.

### Clinical assessment and data-driven decomposition of psychopathology

The clinical assessment consisted of a computerized and modified version of the neuropsychiatric interview Kiddie-Schedule for Affective Disorders and Schizophrenia (K-SADS). This interview assesses psychological treatment history and lifetime occurrence of psychopathology. For participants aged 11-17, information from collaterals were included, while this was the only source for 8-10 year old participants (40). Attempting to separate domains and uncover the empirical correlation structure of psychopathology across diagnostic boundaries, using the larger set of participants (n=6487), we submitted 129 clinical symptom score items, covering 18 clinical domains, to ICA using ICASSO (45), as reported previously (19). The analysis yielded seven psychopathology components. In addition, as a general measure of psychopathology, we computed the mean subject weight across these components.

### Cognitive assessment and definition of a general cognitive factor

Participants completed a computerized test battery evaluating a range of cognitive domains (43). Retrieving a single factor to estimate general cognitive ability, conceptually similar to the IQ score, we included performance scores from 17 diverse tests in a principal component analysis (PCA) without rotation in the larger set of participants (n=6487) as reported previously (19). Due to high correlations between test scores and age, all raw scores were age residualized by linear regression and standardized before running the PCA. The first factor was extracted as a general measure of cognitive function (gF), explaining 12% of the variance.

### MRI acquisition and processing

MRI scans were acquired on a single 3T Siemens TIM Trio whole-body scanner without any software or hardware upgrades (40) (see Supplemental Methods). We used FreeSurfer 5.3 (http://surfer.nmr.mgh.harvard.edu) for cortical reconstruction and volumetric segmentation (46-49). To assess the quality of the cortical reconstructions, we implemented a flagging procedure based on robust PCA for detecting signal to noise-, and segmentation outliers (50, 51). We carefully inspected flagged datasets, and minor edits were performed when necessary (n= 459). Scans from 63 participants were excluded due to poor image quality and had poor Euler number (52) see Supplemental Table S2.

Based on previous implementations (37, 53, 54), for each participant, we sampled signal intensities from the non-uniform intensity normalized volume (nu.mgz) using the FreeSurfer function mri_vol2surf. For each vertex, GM intensities were sampled at six equally spaced points, starting from the gray/white boundary and ending 60% into cortical GM. We selected this endpoint to minimize contamination of voxels containing cerebrospinal fluid. WM intensities were sampled at each vertex at 10 equally spaced points, starting from the gray/white boundary and ending at a fixed distance of 1.5 mm into subcortical WM. To obtain single separate measures of GM and WM intensity per vertex, we calculated the average intensity value for each tissue type. GWC was then computed as: 100 * (white – gray)/[(white + gray)/2] (37), such that a higher value reflects greater difference between GM and WM signal intensities. The GWC surface maps were smoothed using a Gaussian kernel of 10 mm full width at half maximum.

We then performed ICA to decompose the GWC surfaces into spatially independent modes of variations using ICASSO (45). We used a model order of 15, pragmatically chosen based on a compromise between data reduction and total explained variance.

We refer to Supplemental Methods for processing procedures for comparison between GWC and the complimentary method of T1/T2 ratio.

### Statistical analyses

We first tested the associations between each GWC IC and age by computing the T statistics of the linear and quadratic age effects on each GWC IC using General linear models (GLMs), covarying for sex, and p-values were corrected using the false discovery rate (FDR) procedure with a significance threshold of 0.05 (55). To visualize the observed effects we used locally weighted scatterplot smoothing (loess) and ggplot2 (56) in R (https://www.r-project.org/). We also used loess to assess WM and GM separately (see Supplemental Methods).

Second, we tested for associations between the psychopathology component loadings and gF and GWC ICs using GLMs with age and sex as covariates. We also report associations with the quadratic age term when significant. For each of the 15 GWC ICs, we ran GLMs where either one of the 7 psychopathology ICs, mean psychopathology score or gF was included as an independent variable, resulting in 135 (15*9) models. The p-values obtained from all models were then corrected using the FDR procedure with a significance threshold of 0.05 (55). For follow-up analyses concerning brain size and ethnicity see Supplemental Methods. To investigate possible interactions between age and psychopathology and gF on GWC ICs, we repeated the above models including an additional interaction term between these variables. To assess the dynamics of effects not necessarily well modeled by GLM we also performed a sliding window analysis and a subgroup visualization (see Supplemental Methods).

Third, to allow for comparison with a more conventional vertex wise approach, we examined the vertex wise effects of age, covarying for sex, and for psychopathology and gF, covarying for sex and age, on GWC, using GLM as implemented in the Permutation Analysis of Linear Models (PALM) toolbox (57). To assess statistical significance we used 10,000 permutations and family wise error (FWE) correction with threshold-free cluster enhancement (TFCE) (58) and a significance threshold of p<0.05, corrected. For follow-up analyses where thickness was added as a vertex wise co-variate see Supplemental Methods.

## Results

### ICA decomposition of GWC

Figure 1 shows the 15 GWC components from ICASSO. In total, these components explained 29% of the total variance. With some notable exceptions, the maps were highly bilateral and symmetrical, and were region specific, except for a single global component. Regional specification corresponded well with regions separated within T1/T2 ratio maps and histological profiles, both concerning myelin content and developmental patterns (23, 24, 33).

**Figure 1.**
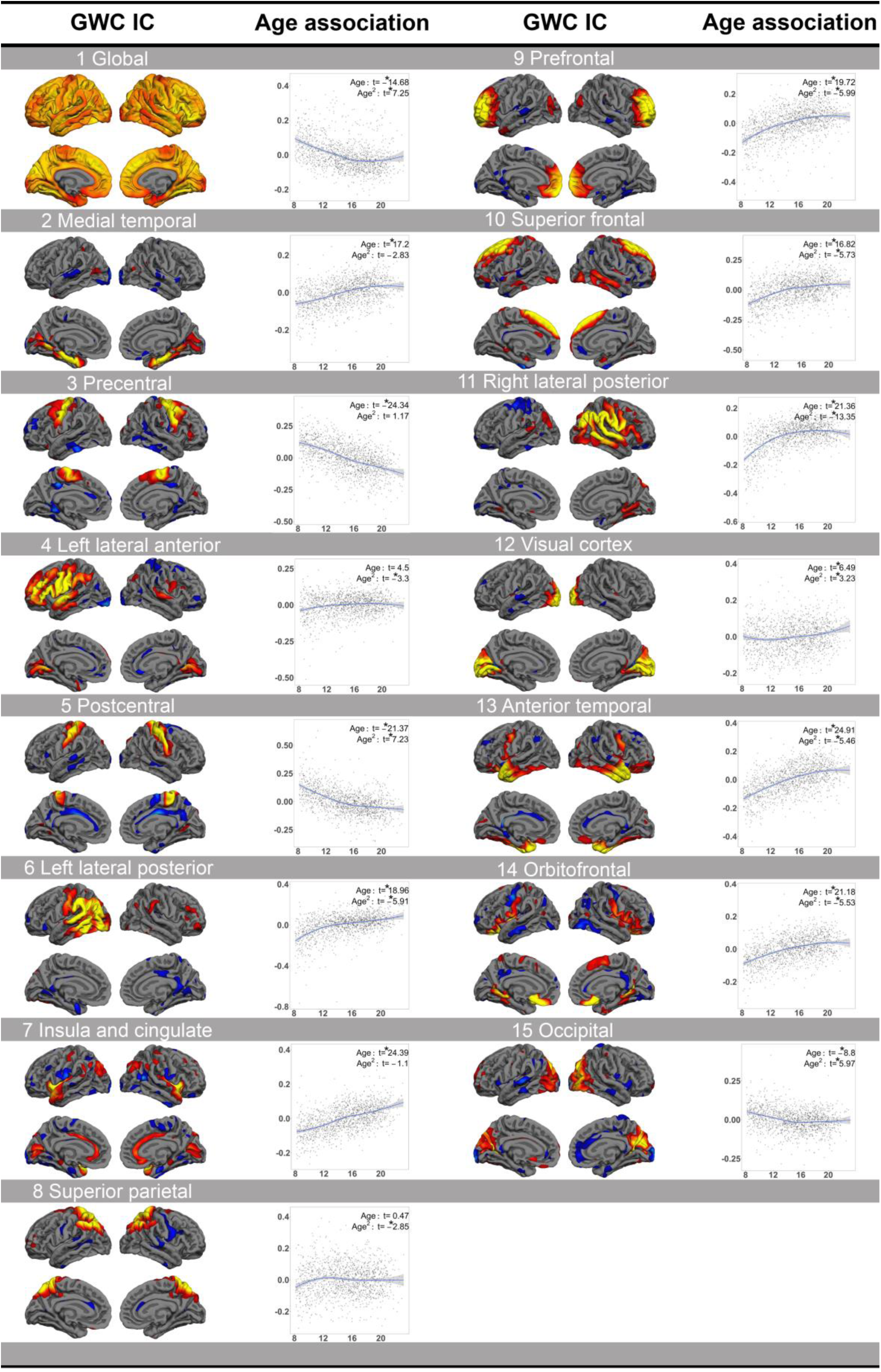
GWC independent components (ICs) and their associations with age. The GWC IC columns show the number, name and anatomical representation. GWC IC maps are thresholded at 1-3 standard deviations (SDs), except “1 Global” which is thresholded at 1-9 SDs. The age association columns show loess visualizations of the age associations. The t statistics for age and age^2^, covarying for sex, are shown in the top right corners. * FDR corrected p <= 0.05

### Associations between GWC and age

The global GWC component showed a marked negative linear age association (t = −14.68, p= 1.27e-45). Importantly, all remaining GWC ICs must be understood as reflecting independent variance beyond this global component. Components reflecting pre- and postcentral cortices showed strong linear negative age-associations (t=-24.34, p=3.70e-110 and t =-21.37, p=1.92e-88 respectively), and the occipital component also showed a highly significant negative association (t=-8.8, p=3.83e-18). The remaining ICs showed either slight or marked linear positive age associations with t-values ranging from 6.49 to 24.91 (all p< 0.05). The superior parietal and left lateral anterior components did not show a significant linear age effect. Loess visualization of the associations between GWC ICs and age are shown in Figure 1. See Supplemental Results- and Figure S1 for loess visualization of GM and WM separately.

### Associations between GWC and psychopathology and gF

Figure 2 shows the results from the GLMs testing for associations between GWC and clinical and cognitive scores. There were several significant associations between GWC and anxiety (IC2) and prodromal psychosis (IC4). Specifically, higher loading on anxiety or prodromal psychosis were both associated with higher GWC in left lateral posterior (t=3.01, p=1.78e-02 and t=2.85, p=2.60e-02, respectively) and insula and cingulate cortices (t=4.55, p=1.33e-04 and t=3.8, p=2.28e-03, respectively), and with lower contrast in pre- (t=-3.51, p=4.86e-03 and t=3.68, p=2.75e-03, respectively) and postcentral cortices (t=-3.28, p=8.91e-03 and t=-2.58, p=4.96e-02, respectively). Anxiety showed additional positive associations in prefrontal (t= 3.47, p=5.27e-03) and right lateral posterior cortices (t=3.00, p=1.78e-02). Prodromal psychosis showed an additional positive association in the visual cortex (t=4.38, p=2.45e-04), and additional negative associations in medial temporal (t=-3.03, p=1.78e-02) and superior frontal cortices (t=-3.16, p=1.29e-02).

**Figure 2.**
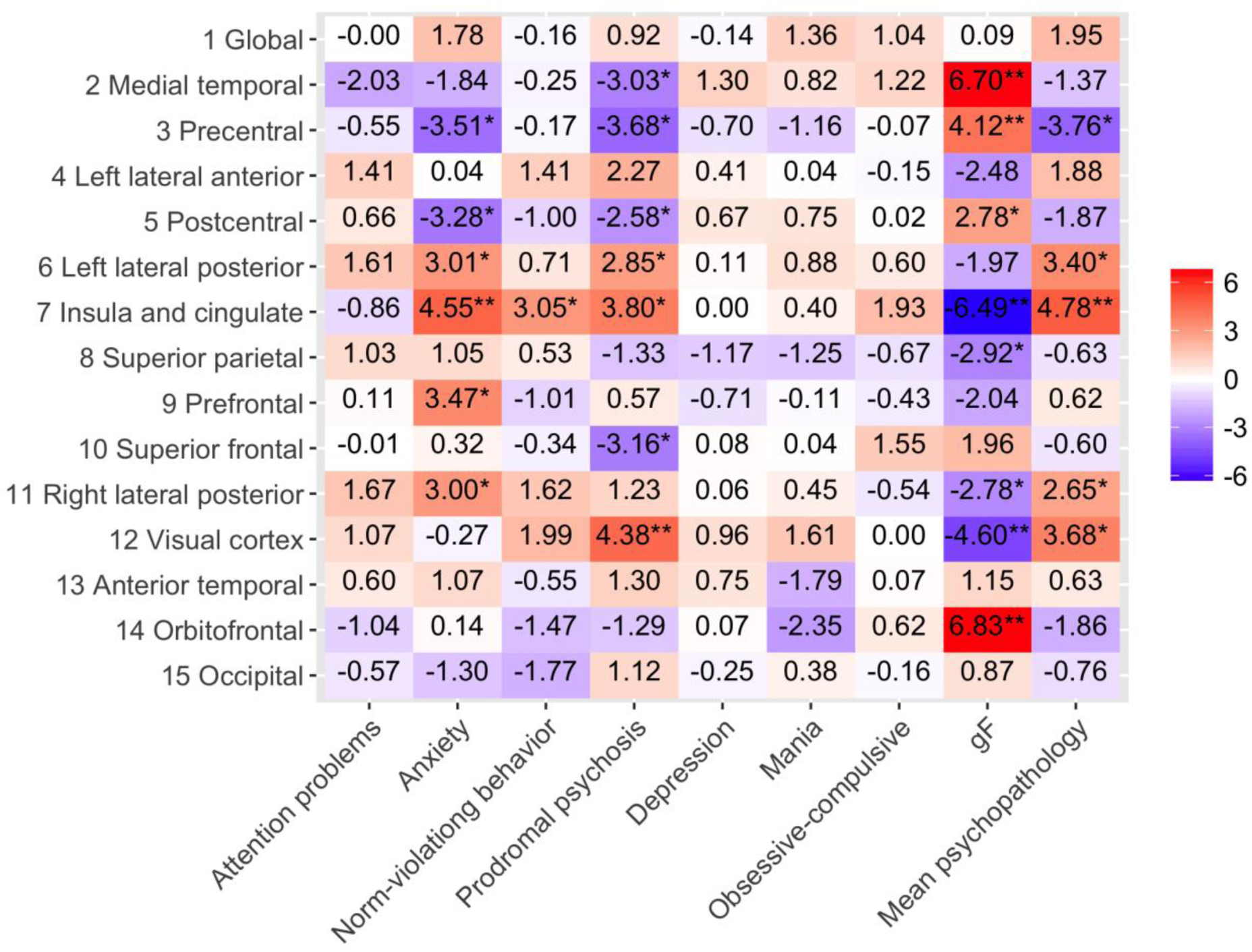
T statistics of the associations between GWC and psychopathology. The y-axis shows each GWC independent component (IC). The x-axis shows each psychopathology IC, gF, and mean psychopathology score. The map is color scaled so that red squares show positive associations, while blue squares show negative associations, covarying for age, age^2^ when significant and sex. * corrected p <= 0.05, ** corrected p <= 0.01.

Except for a single positive association between norm violating behavior and GWC in insula and cingulate cortices (t=3.05, p=1.76e-02), we found no significant associations between the other psychopathology components and GWC. There were, however, significant associations between mean psychopathology score and several GWC components, but these overlapped with the effects of anxiety and/or prodromal psychosis.

General cognitive ability, as indexed by gF, was positively associated with GWC in medial temporal (t=6.70, p=2.09e-09), pre- (t=4.12, p=6.70e-04) and postcentral (t=2.78, p=3.07e-02), and orbitofrontal cortices (t=6.83, p=1.76e-09), and negatively associated with GWC in insula and cingulate (t=-6.49, p= 5.48e-09), superior parietal (t=-2.92, p=2.19e-02), right lateral posterior (t=-2.77, p=3.07e-02), and visual cortices (t=-4.60, p=1.25e-04). Six of these eight associations between general cognitive ability and GWC spatially overlapped with associations between psychopathology and GWC, but with effects in the opposite direction. The correlation with gF loading was −0.14 and −0.13 for anxiety and prodromal psychosis loading respectively. See Supplemental Results for brain size added as a co-variate (Figure S2), and for analyses within the two largest ethnicity groups (Figure S3). Briefly, adding brain size expectedly reduced effect sizes somewhat, but association patters were retained. Contrarily, performing separate analyses within Caucasians and African Americans/Blacks indicated differential association patterns.

We found no significant interactions between age and psychopathology or gF on the GWC ICs. For GLMs with interaction terms, sliding window results, and loess subgroup visualization, we refer to the Supplemental Material (Supplemental Figures S4-6).

### Vertex wise associations between GWC and age, psychopathology and gF

Figure 3 shows the results from the vertex wise analyses. Briefly, permutation testing revealed a near global negative association between age and GWC, indicating lower GWC in widespread regions with higher age. Strongest negative associations were seen in regions around the central sulcus (Supplemental Table 3). These results generally correspond well with the results from the ICA approach. For results where thickness was added as a vertex wise co-variate, see Supplemental Results- and Figure S7. Additionally, for per-vertex full- and partial correlations controlling for age between thickness and GWC see Supplemental Figures S8-9. In short, adding thickness had minor effects on results, and correlations between GWC and thickness were generally low. For results on the vertex wise associations between psychopathology and gF and GWC, see Supplemental Results. Briefly, there were no significant associations of the psychopathology components, but negative significant associations with gF and GWC (Supplemental Figure S10).

**Figure 3.**
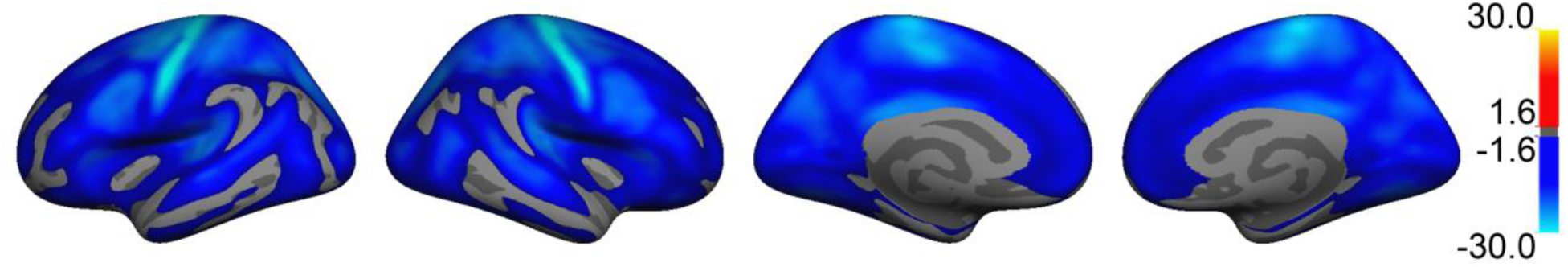
Vertex wise associations between age and GWC. T statistics are masked by FWE corrected p-values thresholded at a minimum –log(p) of 1.6 to correct for two hemispheres. Blue regions represent negative age associations.

### Comparison between GWC map and T1/T2 ratio map

Figure 4 depicts the mean GWC surface map from all participants compared to the Conte69 mean T1/T2 ratio map provided by HCP, (see Supplemental Results for in detail descriptions). For mean GM and WM surfaces displaying intensity spread across the cerebrum we refer to Supplemental Figure S11.

**Figure 4.**
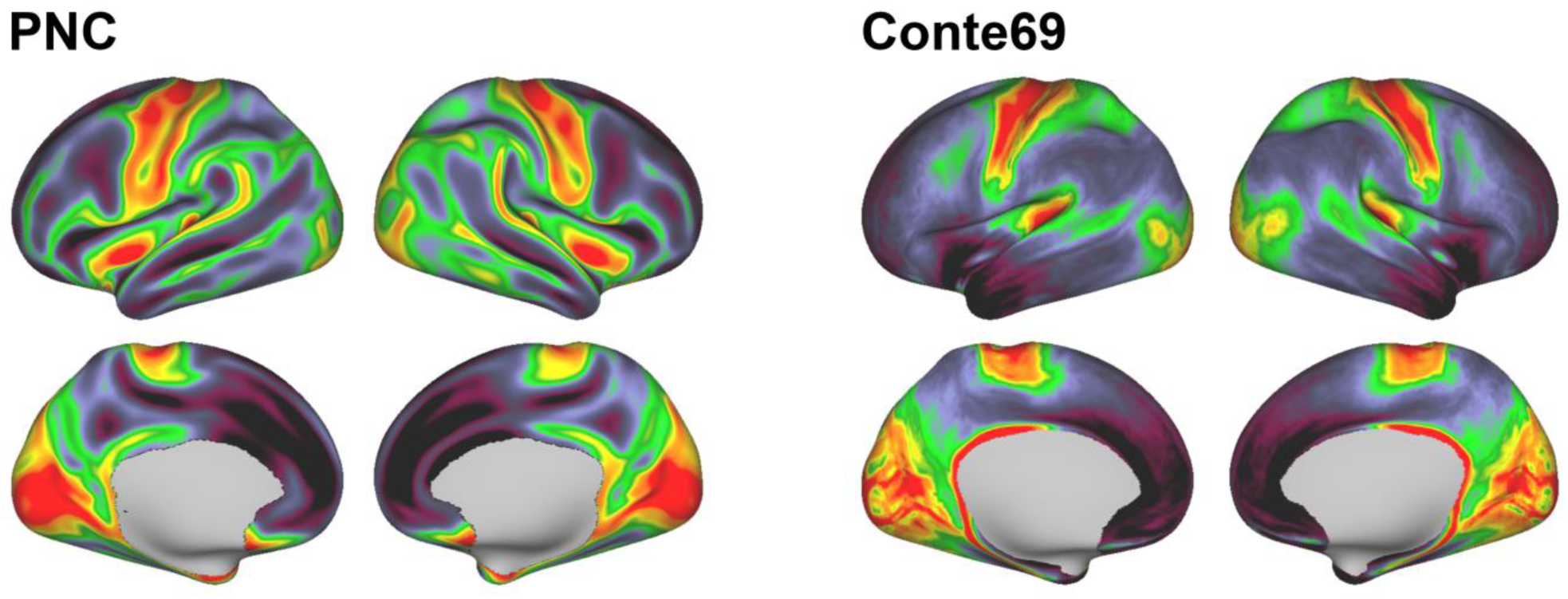
Mean GWC map vs T1/T2 ratio map. PNC depicts the mean GWC surface maps of the current sample. Warm colors represent regions with lower GWC, while cold colors represent regions with higher GWC. Conte69 depicts a mean T1/T2 ratio map from young adults, based on 69 subjects (male=38, female=31, age=9-45) (77). Warm colors are thought to represent regions with a high amount of intracortical myelin, while cold colors represent regions of low myelin content.

## Discussion

Cerebral myeloarchitecture develops across childhood and adolescence and, based on the current study, individual variability in the difference between intracortical and subjacent WM myelination likely has clinical relevance for a range of mental disorders, in particular anxiety and psychotic disorders. Based on T1-weighted brain scans from 1467 children and adolescents, we approximated myelination using the contrast in signal intensity between intracortical and subjacent WM across the brain. We found a global blurring of the GWC at higher ages, possibly in part reflecting protracted intracortical myelination compared to subjacent WM, with additional regional developmental patterns. Across individuals, regional GWC was associated with symptom burden, with significant associations to symptoms of anxiety and prodromal psychosis, and with general cognitive ability.

A broad range of psychopathology components were identified using a data-driven approach and regional GWC was found to be associated with symptom levels of psychopathology within components reflecting several forms of anxiety and a component reflecting positive prodromal and advert psychosis. Higher levels of either anxiety or prodromal psychosis were associated with higher GWC in left lateral posterior and insula and cingulate cortices, and lower contrast in sensorimotor cortices. Anxiety also showed positive associations in prefrontal and right lateral posterior cortices, while prodromal psychosis showed a positive association in the visual cortex and negative associations in medial temporal and superior frontal cortices. Although our original hypothesis was that increased symptom burden would be linked to regionally higher GWC, effects in both directions have been reported in prior studies (see below).

In addition to well documented WM aberrations (19, 59-61), there is accumulating evidence for abnormal intracortical myelination as a potential mechanism for brain network dysfunction in neurodevelopmental disorders, and psychosis in particular (16). Myelin-related abnormalities in psychosis is indicated by genetic association studies (62), post mortem studies (63, 64) and from the notion that psychotropic treatments have effects on myelin, and its plasticity and repair (65). Two previous studies have investigated GWC in adults with schizophrenia with similar methods. Kong et al. (38) reported decreased GWC in frontal and temporal regions, while Jørgensen et al. (37) in a larger study reported the opposite i.e. increased GWC, yet in separate sensorimotor regions. Somewhat inconsistent with these studies, our results indicate that higher psychosis symptom levels in adolescents are associated with decreased GWC in pre- and postcentral cortices extending into superior frontal cortex and with increased GWC in left lateral posterior, insula and cingulate, and visual cortices. The discrepancy could be due to analysis approach, sample age, or clinical characteristics. In addition to associations with prodromal psychosis, we also found associations with anxiety in partly overlapping regions.

We also investigated associations between general cognitive ability and GWC, hypothesizing that higher cognitive ability would be associated with lower GWC. The results indeed showed negative correlation in insula and cingulate, superior parietal, right lateral posterior, and visual cortices, but also positive associations in medial temporal, pre- and postcentral, and orbitofrontal cortices. Interestingly, most of these regions overlapped with regions in which we found associations between GWC and psychopathology, with effects consistently in the opposite direction.

A few prior studies measuring either GWC or T1/T2 ratio have reported associations with cognitive functioning (25, 26, 66). For example, a study on youth reported that brain age gap computed using GWC was related to IQ (25). The effects in this study were driven by “decreased age” in sensorimotor regions and “increased age” in association cortices. Correspondingly, we found higher GWC in sensorimotor regions for participants with high cognitive functioning, and lower GWC in large posterior regions, particularly in the right hemisphere. The observation that several of the GWC regions associated with gF overlapped with regions associated with psychopathology, correspond well with a separate study, also on the PNC sample, which reported that gF and mean psychopathology score share a genetic contribution (19).

In our follow-up analyses within Caucasians and African Americans/Blacks separately, there were discrepancies in the association pattern between GWC and both psychopathology and gF. An exploration of possible genetic or environmental factors accounting for these discrepancies lie beyond the scope of the current study. However, we note that neighbourhood crime rate and parental education has previously been found to partially mediate the association between ethnicity and cognitive function in the PNC sample (67, 68).

The global GWC component showed an age-related decrease across late childhood and adolescence. Although the microstructural underpinnings represent changes both within GM and WM, our results do fit well with prior studies employing cross-sectional MRI cortical intensity measures or post mortem histology of intracortical myelin. For instance, a study investigating cortical signal intensity reported a steady age-related decrease from childhood to adulthood (53), while a study employing T1/T2 ratio reported what was interpreted as an ongoing intracortical myelination process from 8 years of age extending well into adulthood (66). Correspondingly, a post mortem study reported that the maturation of human intracortical myelination extends past late adolescence (29).

Beyond the global age-related decrease in GWC, regional components showed both positive and negative age effects, which could in sum reflect regionally protracted and accelerated development, respectively. For instance, components encompassing primary sensorimotor or occipital regions showed negative associations with age. These regions have a high myelin content and mature early (23, 24). In comparison, e.g. the frontal lobe and temporal pole are known to continue intracortical myelination considerably longer (66, 69), and the frontal lobe shows peak gray matter signal intensity later than the age range of the current sample (53). Components capturing these regions in the present study showed positive age associations, which together with the global negative age effect, might indicate regionally protracted development.

Our GWC map showed high consistency with a mean T1/T2 ratio map provided by HCP (24). The moderate correlation between the two maps could partly reflect differences in age range, and scanner related properties. Still, the strength of the correlations indicates that although the measures share variance, they also have specific, unique properties.

A limitation of the current study is that age-related differences in GWC cannot be specifically attributed to protracted intracortical myelination. Although our results converge with prior histological- and MRI studies, the microstructural underpinnings of the measure are likely highly complex. Logically, GWC is a combination of both GM and WM, and loess visualizations indeed indicated similar age trajectories. Additionally, the biological interpretation of T1/T2 ratio as a proxy for intracortical myelin has come under scrutiny (70, 71). Moreover, the intensity in T1-weighted images also reflects other biological properties, such as water content, iron, and dendrite density (39, 72, 73). Another study limitation is the potential for partial volume effects due to sampling intensities close to the gray/white boundary. We chose this approach to include deeper, more myelinated layers of cortex. Partial volume effects could vary depending on cortical thickness, although vertex wise correlations between GWC and thickness were generally low, and adding thickness as a regressor had minor effects on vertex wise results. Lastly, the cross sectional design, which is a non-optimal indirect way of studying development (74) is a limitation to the current study.

The relevance of the current study is twofold. First, studies of individuals at risk for mental disorders have typically used genetic risk from family history, or clinical risk measured as symptoms in help seeking individuals (75). In contrast, the PNC sampling strategy taps into the continuum of psychopathology in youth. Thus, our findings may help define markers of neural dysfunction at both an earlier age and stage of psychopathology development. This is critical for identifying biomarkers useful for early detection, and eventually for prevention and early intervention (75, 76). The finding that aberrations are already present in generally healthy youth with increased levels of anxiety and prodromal psychosis is an important addition to the literature. Second, we used data-driven approaches to identify both a broad range of psychopathology components and regional patterns in GWC, aiming to improve sensitivity and specificity, as associations between brain structure and psychopathology are typically small or moderate (see e.g. (51)).

To conclude, the results of the current study showed that GWC globally decreases across late childhood and adolescence, likely partly reflecting the extended maturation of intracortical myelination as compared to subjacent WM. We additionally found regional developmental patterns possibly reflecting accelerated and protracted myelination. Across individuals, regional GWC was associated with symptom burden of anxiety and prodromal psychosis and general cognitive ability, supporting the clinical and neurocognitive relevance of this tissue contrast measure.

## Acknowledgments and disclosures

This work was supported by the Department of Psychology, University of Oslo (LBN and CKT), the Research Council of Norway (#223273, #249795, #230345), the South-Eastern Norway Regional Health Authority (#2014097, #2016083), and the European Commission’s 7th Framework Programme (#602450, IMAGEMEND). Support for the collection of The Philadelphia Neurodevelopment Cohort was provided by grant RC2MH089983 to Raquel Gur, and RC2MH089924 to Hakon Hakonarson. The participants were recruited through the Center for Applied Genomics at The Children’s Hospital in Philadelphia. Data for this article was in part collected and shared by the Cambridge Centre for Ageing and Neuroscience (CamCAN). The UK Biotechnology and Biological Sciences Research Council (BB/H008217/1) provided CamCAN funding, with support from the UK Medical Research Council and University of Cambridge, UK. Data were also in part provided by the Human Connectome Project, WU-Minn Consortium (Principal Investigators: David Van Essen and Kamil Ugurbil; 1U54MH091657) funded by the 16 NIH Institutes and Centers that support the NIH Blueprint for Neuroscience Research; and by the McDonnell Center for Systems Neuroscience at Washington University. Parts of the data in the present paper have previously been presented as a poster at the Society for Neuroscience 2017 annual meeting, and a preprint is published on bioRxiv.org. The authors report no biomedical financial interests or potential conflicts of interest.

## Supplemental information

### Supplemental Methods and Materials

#### Participants

The institutional review boards of the University of Pennsylvania and the Children’s Hospital of Philadelphia (CHOP) approved all study procedures, and written informed consent, as well as parental permission for individuals under the age of 18 years, were obtained (1). CHOP recruited participants through a previous separate study enrollment (2). 9498 participants completed psychiatric and cognitive assessment, and 1601 of these individuals were randomly selected for neuroimaging after stratification by age and sex (3).

#### MRI acquisition

PNC: Signal excitation and reception was obtained using a quadrature body coil for transmit and a 32-channel head receiver coil. Gradient performance was 45mT/m, with a maximum slew rate of 200 T/m/s (4). T1 weighted imaging was obtained using a magnetization prepared rapid-acquisition gradient-echo (MPRAGE) sequence (TR=1810 ms; TE=3.51 ms; FoV=180×240 mm; Resolution=0.94×0.94×1.0 mm). Receive coil (i.e. B1) shading was reduced by selecting the Siemens pre-scan normalize option, which corrects for B1 inhomogeneity based on a body coil reference scan (4).

Cam-CAN: MRI scans were aquired on a 3T Siemens TIM Trio System, and signal excitation and reception was obtained using a 32 channel head coil. High resolution 3D T1-weighted structural images were obtained using a Magnetization Prepared Rapid Gradient Echo (MPRAGE) sequence with the following parameters: Repetition Time (TR) =2250 milleseconds; Echo Time (TE) =2.99 milliseconds; Inversion Time (TI) =900 milliseconds; flip angle =9 degrees; field of view (FOV) =256mm x 240mm x 192mm; voxel size =1mm isotropic; GRAPPA acceleration factor =2; acquisition time of 4 minutes and 32 seconds (5).

TOP: MRI scans were acquired on a single 3T GE 750 scanner. Signal excitation and reception was obtained using a 32-channel head receiver coil. T1 weighted imaging was obtained in the sagittal plane, with 1mm isotropic voxels and pulse sequence “Brain Volume imaging”

(BRAVO). Flip Angle: 12, TI: 450, Receiver Bandwidth: 31.25, Freq: 256, Phase: 256, Freq DIR: S/I, NEX: 1.00, Phase FOV: 1.00.

#### MRI processing of GWC and T1/T2 ratio

To compare the gray/white matter contrast (GWC) measure with the complimentary method of T1/T2 ratio, we created a mean GWC surface map of all participants, which was then converted to gifti format and transformed into the Human Connectome Project (HCP, (6)) standard mesh of 164,000 vertices (7). We then performed vertex wise Pearson correlations of our GWC mean map and the Conte69 mean T1/T2 ratio map provided by the HCP (8). To rule out confounding effects induced when comparing maps from different samples, we also created both GWC and T1/T2 ratio (8, 9) mean surface maps as reported, but from the same subjects in two independent samples; Cam-CAN, obtained from the CamCAN repository (available at http://www.mrc-cbu.cam.ac.uk/datasets/camcan/) (5, 10), (male=10, female=8, age=18-29), and TOP (11, 12) (for MRI protocols, see above), (male=10, female=10, age=15-54). This allowed us to correlate maps obtained from the same subject using the two different methods.

#### Statistical analyses

##### Loess visualization of gray- and white matter associations with age

In order to investigate the differential gray and white matter (WM) contributions to the observed associations between GWC and age, we performed a follow-up visualization of the association with age for each matter type separately. It should be noted that raw signal intensities are not constant across participants for the same underlying relaxation parameters due to session- and participant-specific optimization procedures performed during MRI acquisition. Thus, signal intensities should be normalized to correct for inter-subject variations in scaling factors. Intensities were first normalized by ventricular CSF or voxel intensities outside of the scull. Both these metrics were strongly associated with age and using these for normalization would therefore unduly affect the results of subsequent analyses. We therefore performed a loess visualization of raw signal intensity scores sampled within different gray matter (GM) and WM regions, transformed to z-scores and plotted as a function of age.

##### Main effects model with brain size as a co-variate

In order to correct for possible effects of brain size in our original main effects model on the associations of GWC ICs and psychopathology and gF, we performed a follow-up analysis. We included as an additional co-variate a variable named “brainSegVolnotVent”, a Freesurfer output including all voxels from aparc and aseg.mgz that are neither background nor brain stem, and excludes ventricles, CSF, and choroid plexus.

##### Main effects model within Caucasians and African Americans/Blacks separately

In order to assess whether the associations of GWC ICs and psychopathology and gF were different within the two largest ethnicity groups, we re-ran the original main effects model within individuals reporting that they were Caucasian only (n=680) or African American/Black only (n=635).

##### Sliding window analysis

In order to investigate dynamics of possible effects not necessarily well modeled by standard linear models, we performed GLM sliding window analyses, with width of 5 years and steps of 1 year. Here, GWC ICs were the dependent variable, age and sex were covariates, and only psychopathology ICs or the gF component that had already shown a main effect, were tested as separate independent variables. At each step, the t-statistics representing the association between relevant psychopathology ICs or gF on GWC ICs were extracted and standardized using Cohen’s *d*. All p-values were FDR corrected with a significance threshold of 0.05 (13) for all windows.

##### Loess subgroup visualization

To visually assess the effects found by GLM and sliding window analyses we performed a subgroup visualization by loess ggplot in R. Again, only psychopathology ICs or the gF component that had already shown a main effect, were tested as separate independent variables. We fitted a curve on the full sample for the association between GWC ICs and age. We then identified participants with the lowest loading on relevant independent variables, and repeated the same spline fitting procedure on sub groups, after removing 50%, 80% and 90% of these participants respectively. Thus, beginning with the full sample and then continuously removing the “healthiest” (or for gF “lowest” performing) individuals, in order to inspect changes in the association. We chose the above-described four percentage groups after reviewing the spread of each relevant independent variable. Splines were then bootstrapped 1000 times in order to extract confidence intervals.

##### PALM analyses with thickness as a per vertex co-variate

In order to investigate possible effects of thickness on our GWC vertex wise analyses, we re-ran PALM analysis for those models showing significant associations, specifically the associations between GWC and age, and GWC and gF. We now added thickness as a vertex wise co-variate, and performed only one permutation. We then compared uncorrected t-statistics surface maps.

### Supplemental results

#### Loess visualization of gray- and white matter associations with age

We plotted z-scores of raw signal intensities sampled within different GM and WM regions as a function of age (see Supplemental Figure S1). Although these results should be viewed with caution as they are not statistical analyses and on raw signal intensities, the two tissue types showed overall similar age-associations.

#### Main effects model with brain size as a co-variate

Adding brain size as a co-variate expectedly reduced the effect sizes somewhat, but association patters were largely retained (see Supplemental Figure S2).

#### Main effects model within Caucasians and African Americans/blacks separately

Comparing the t statistic maps of the association between GWC ICs and psychopathology and gF separately within Caucasians and African Americans/Blacks, indicated differential association patterns (see Supplemental Figure S3).

#### Sliding window

Anxiety and prodromal psychosis showed positive age interactions with three separate GWC ICs, while gF showed one positive age interaction. Prodromal psychosis and gF additionally showed three separate negative age interactions (see Supplemental Figure S5). In sum, although we did not find any significant GLM effects with an age interaction term, the sliding window approach still yielded trends toward age interaction effects for anxiety, prodromal psychosis and gF on several GWC ICs. As correction for multiple testing was solely performed across windows and not across the full set of tests, results should be interpreted with caution.

#### Loess subgroup visualization

For the loess subgroup visualization plots of anxiety, prodromal psychosis and gF, see Supplemental Figure S6. In short, the visualizations corresponded well with clinical main effects reported from GLM analyses, as well as the sliding window trends.

#### PALM analyses with thickness as a per vertex co-variate

Comparing uncorrected t statistic maps with thickness added as a vertex wise co-variate to our original results, show highly similar patterns, see Figure S7.

#### Vertex wise associations between anxiety, prodromal psychosis, gF and GWC

Permutation testing revealed no significant associations for either of the psychopathology ICs, but unthresholded t-statistic maps were in general in agreement with the effects revealed through ICA (Supplemental Figure S10). There were vertex wise significant negative associations between gF and GWC in medial and posterior regions of the brain, consistent with the ICA results, albeit not capturing the positive effects.

#### T1/T2 ratio and GWC

Figure 4 depicts the mean GWC surface map from all participants compared to the Conte69 mean T1/T2 ratio map provided by HCP. What is presented as highly myelinated cortical regions within the ratio map, such as sensorimotor and early association regions, showed low GWC, while regions such as the prefrontal and premotor cortex, where myelination is presented as low, showed high GWC. The vertex wise correlation between the two maps was 0.58. We performed vertex wise Pearson correlations of the GWC and T1/T2 ratio mean maps created from the two independent datasets, namely Cam-CAN and TOP. We found vertex wise correlations of −0.54 and −0.77, respectively.

## Supplemental Tables

**Table S1.**
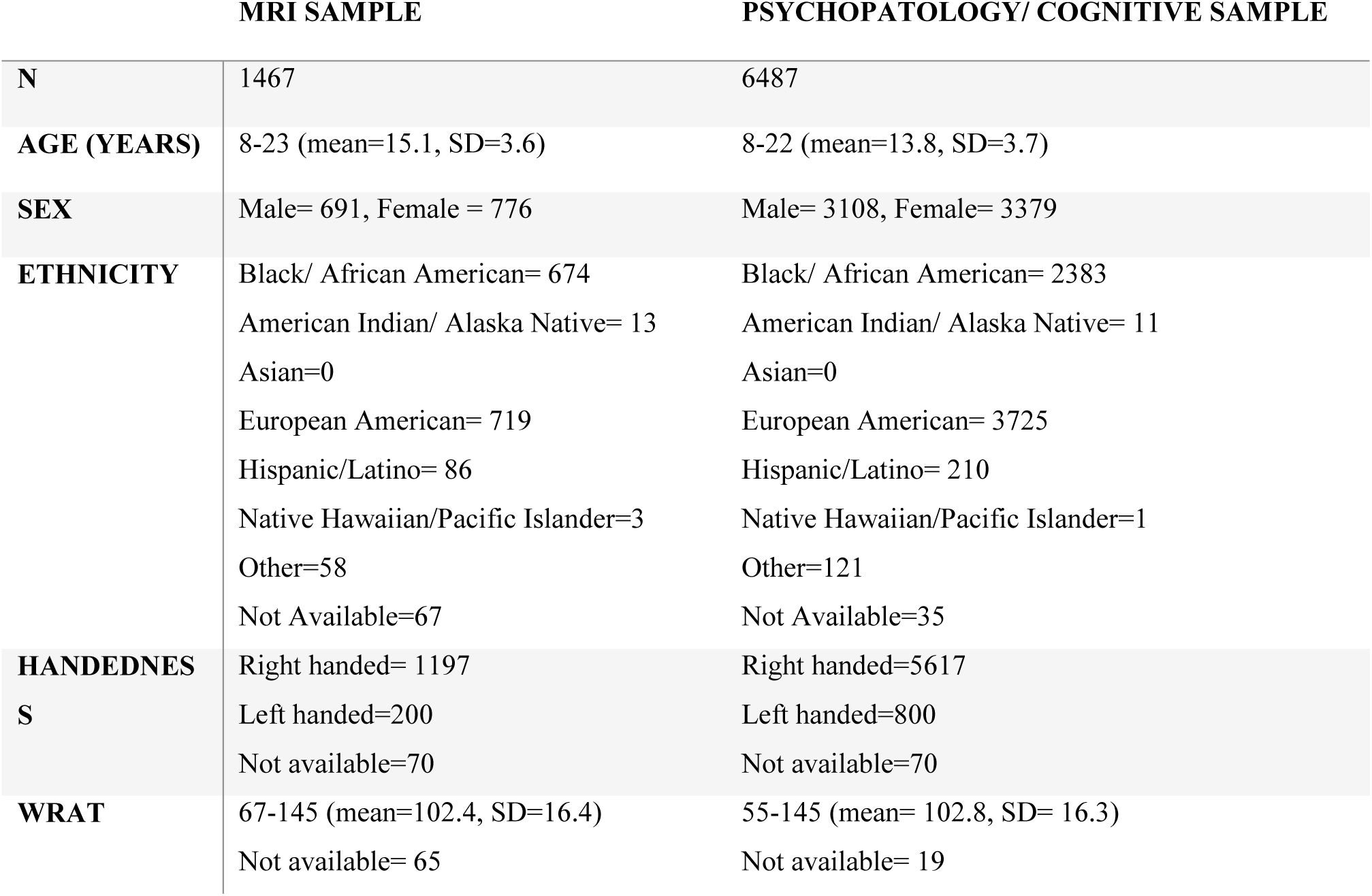
Demographics table of included samples. The total ethnicity numbers exceed the total sample size as several subjects identified as belonging to more than one ethnicity group. WRAT: Wide Range Achievement Test total standard score.

**Table S2.**
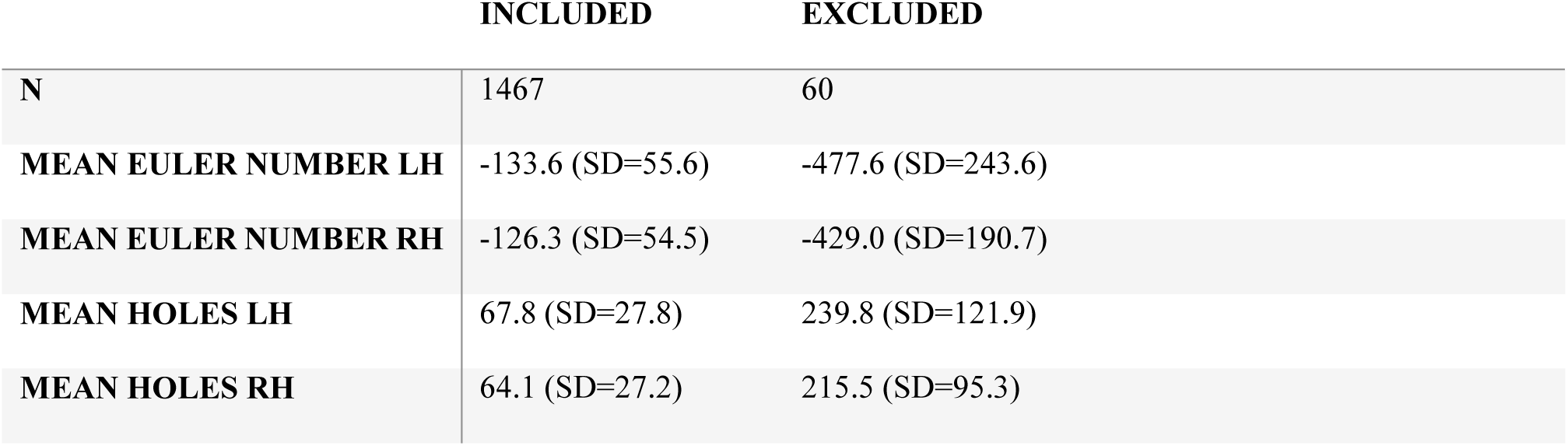
Euler number of excluded (solely due to poor/corrupt image quality) and included subjects. FreeSurfer processing failed for the right hemisphere for 3 subjects due to poor/corrupt image quality, and these were removed from the means of the right hemisphere. Additionally, 4 subjects had missing or too corrupt MRI data to perform Freesurfer processing.

**Table S3.** ROI t statistic overview of age effects. The left column lists the regions of interest. The next two columns depict the mean t statistic in the left and right hemisphere respectively for the effect of age, covarying for sex.

| **STRUCTURAL ROI** | **LH** | **RH** |
| --- | --- | --- |
| **BANKS SUPERIOR TEMPORAL SULCUS** | -9.71 | -11.75 |
| **CAUDAL ANTERIOR CINGULATE** | -8.15 | -7.27 |
| **CAUDAL MIDDLE FRONTAL** | -14.40 | -13.56 |
| **CUNEUS** | -11.23 | -10.66 |
| **ENTORHINA** | -0.72 | -0.53 |
| **FUSIFORM** | -6.43 | -6.25 |
| **INFERIOR PARIETAL** | -6.30 | -7.57 |
| **INFERIOR TEMPORAL** | -4.13 | -2.53 |
| **ISTHMUS CINGULATE** | -12.41 | -11.84 |
| **LATERAL OCCIPITAL** | -7.78 | -8.94 |
| **LATERAL ORBITOFRONTAL** | -5.11 | -5.17 |
| **LINGUAL** | -8.14 | -8.68 |
| **MEDIAL ORBITOFRONTAL** | -2.06 | -2.19 |
| **MIDDLE TEMPORAL** | -2.43 | -2.01 |
| **PARAHIPPOCAMPAL** | -0.61 | -1.53 |
| **PARACENTRAL** | -20.73 | -20.20 |
| **PARSOPERCULARIS** | -10.87 | -11.56 |
| **PARSORBITALIS** | -1.97 | -5.75 |
| **PARSTRIANGULARIS** | -9.60 | -10.20 |
| **PERICALCARINE** | -9.86 | -9.87 |
| **POSTCENTRAL** | -14.51 | -14.51 |
| **POSTERIOR CINGULATE** | -11.34 | -11.11 |
| **PRECENTRAL** | -19.59 | -19.95 |
| **PRECUNEUS** | -13.29 | -12.85 |
| **ROSTRAL ANTERIOR CINGULATE** | -3.74 | -3.66 |
| **ROSTRAL MIDDLE FRONTAL** | -4.81 | -6.33 |
| **SUPERIOR FRONTAL** | -9.44 | -8.61 |
| **SUPERIOR PARIETAL** | -13.95 | -14.76 |
| **SUPERIOR TEMPORAL** | -6.56 | -6.07 |
| **SUPRAMARGINAL** | -7.46 | -9.22 |
| **FRONTAL POLE** | -1.62 | 0.03 |
| **TEMPORAL POLE** | 0.67 | 2.72 |
| **TRANSVERSE TEMPORAL** | -15.28 | -15.85 |
| **INSULA** | -5.47 | -5.23 |

## Supplemental Figures

**Figure S1.**
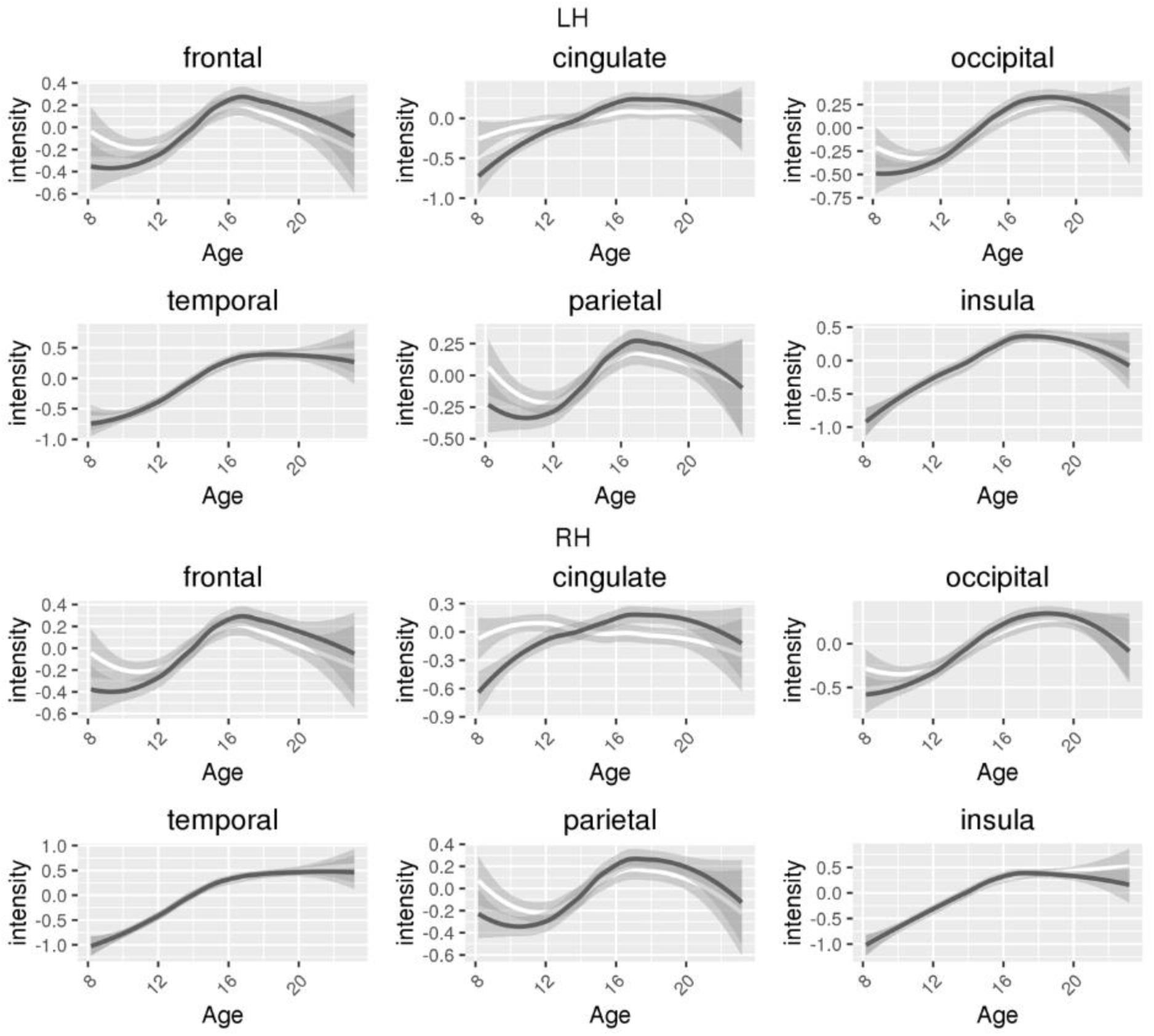
Gray matter (GM) and white matter (WM) z-scores loess visualization. The plots show the fitted curve for raw GM intensity z-scores indicated in dark gray, and raw WM intensity z-scores indicated in white, per cortical lobe. The shaded gray areas represent the confidence interval.

**Figure S2.**
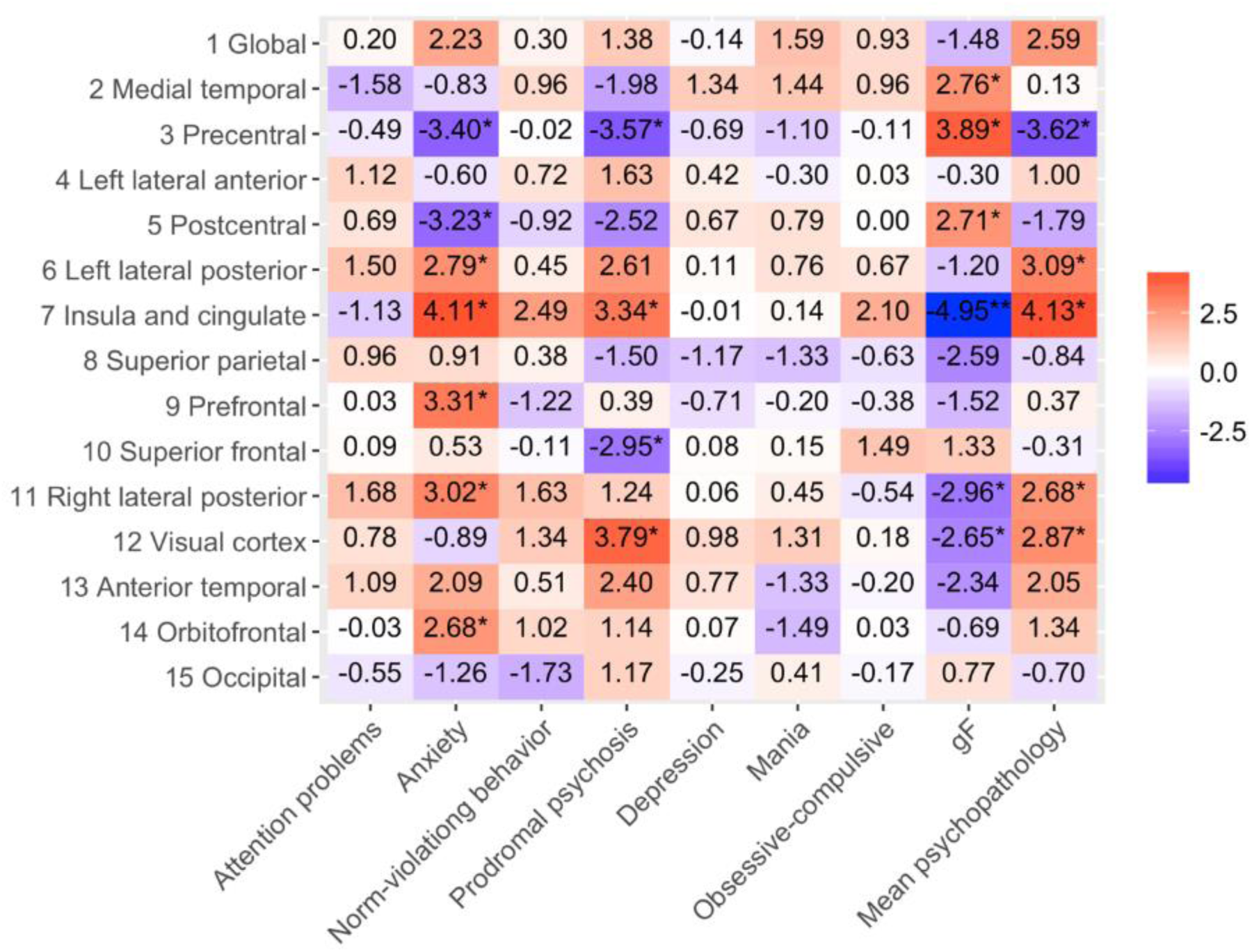
T statistics of the associations between GWC and psychopathology and gF, with brain size as an additional co-variate together with age, age2 when significant and sex. The y-axis shows each GWC independent component (IC). The x-axis shows each psychopathology IC, gF, and mean psychopathology score. The map is color scaled so that red squares show positive associations, while blue squares show negative associations. * corrected p <= 0.05, ** corrected p <= 0.01.

**Figure S3.**
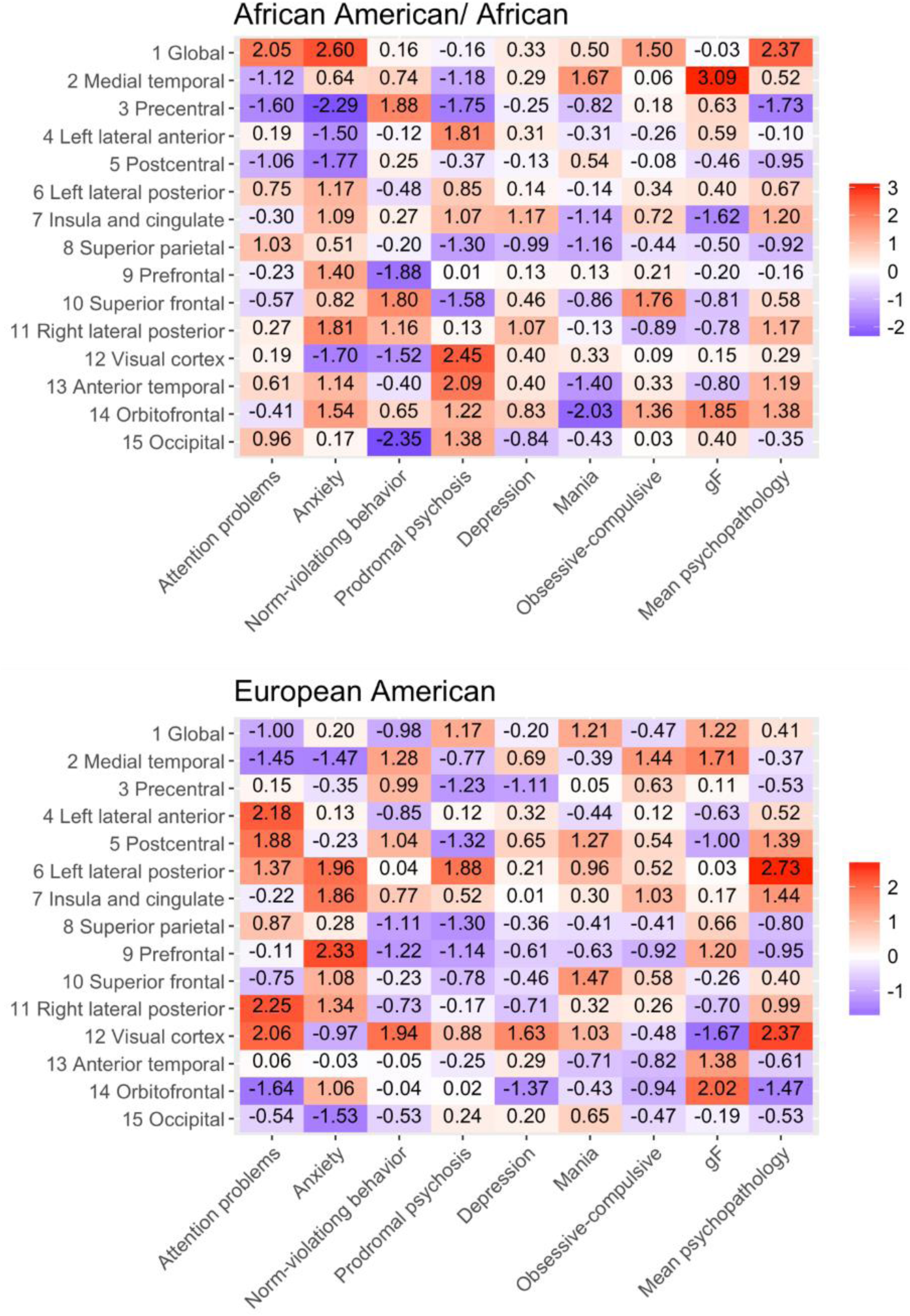
T statistics of the associations between GWC and psychopathology and gF within Caucasians and African Americans/Blacks separately. The y-axis shows each GWC independent component (IC). The x-axis shows each psychopathology IC, gF, and mean psychopathology score. The map is color scaled so that red squares show positive associations, while blue squares show negative associations, covarying for age, age^2^when significant and sex. * corrected p <= 0.05, ** corrected p <= 0.01.

**Figure S4.**
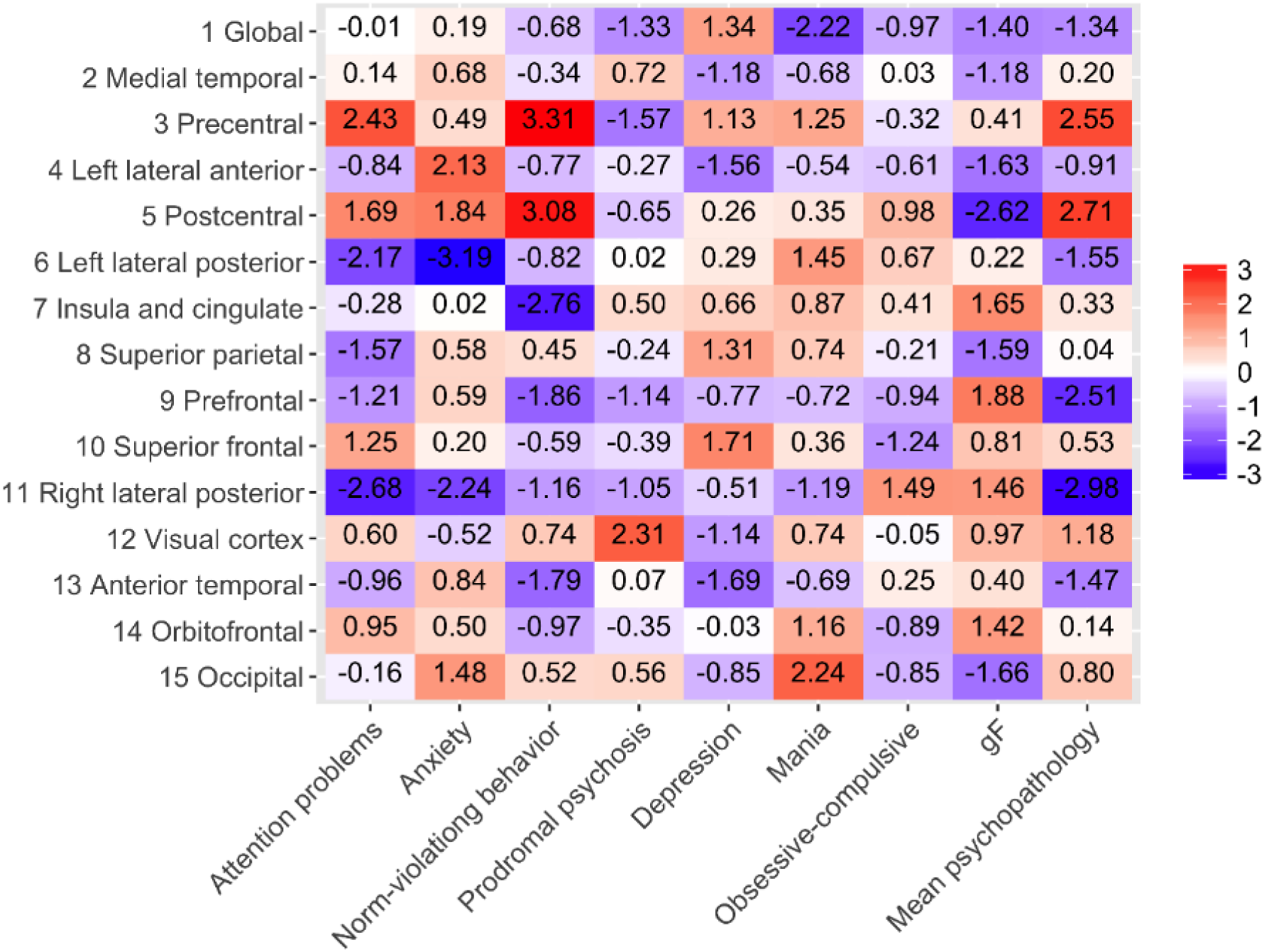
Interaction effects of psychopathology, gF and age on GWC. The y-axis shows each GWC independent components (ICs). The x-axis shows each psychopathology IC, gF and mean psychopathology score. The map is color scaled so that red squares show positive associations, while blue squares show negative associations, covarying for age, age^2^ when significant, and sex. * corrected p <= 0.05, ** corrected p <= 0.01.

**Figure S5.**
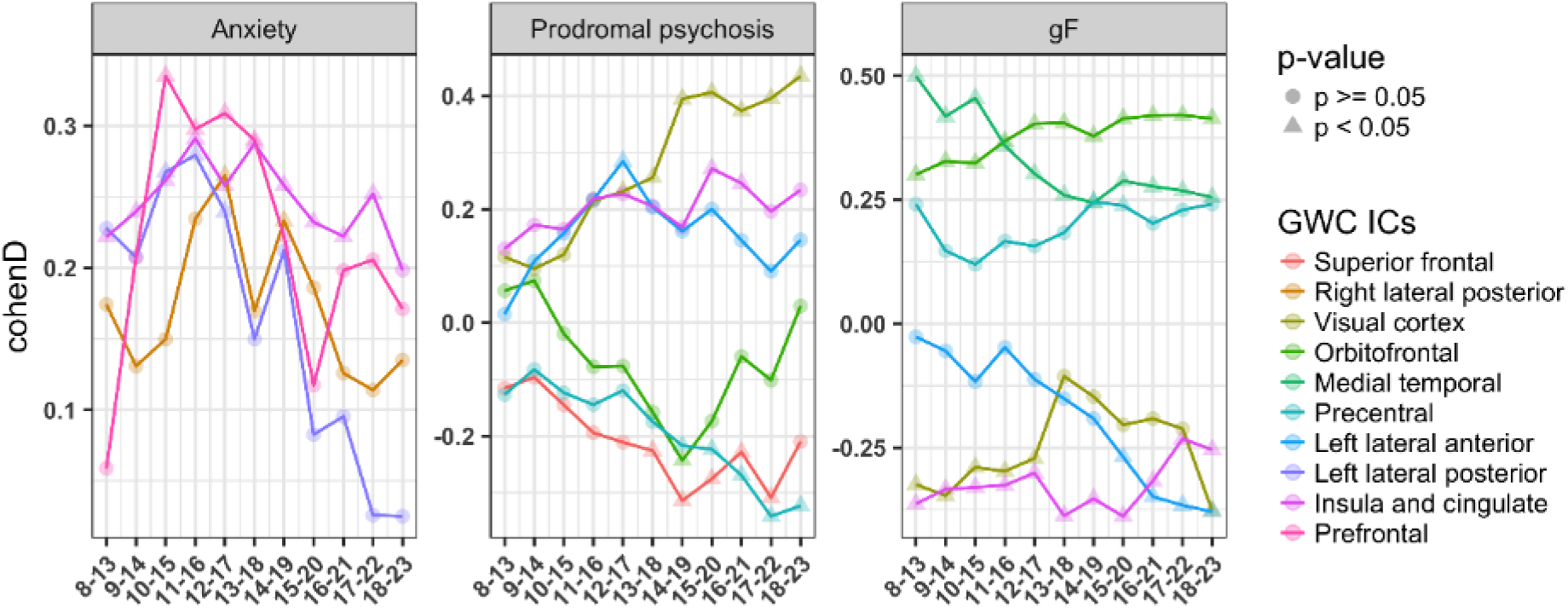
Sliding window analyses, with width of 5 years and steps of 1 year. GWC independent components (ICs) are depicted only if at least one significant age window was found. Left plot shows anxiety, middle plot shows prodromal psychosis and the right plot shows gF. The y-axis represents Cohen’s D value of the strength of the associations, while the x-axis represents the age windows. Triangles on the line represent Cohen’s D values with corrected p < 0.05.

**Figure S6.**
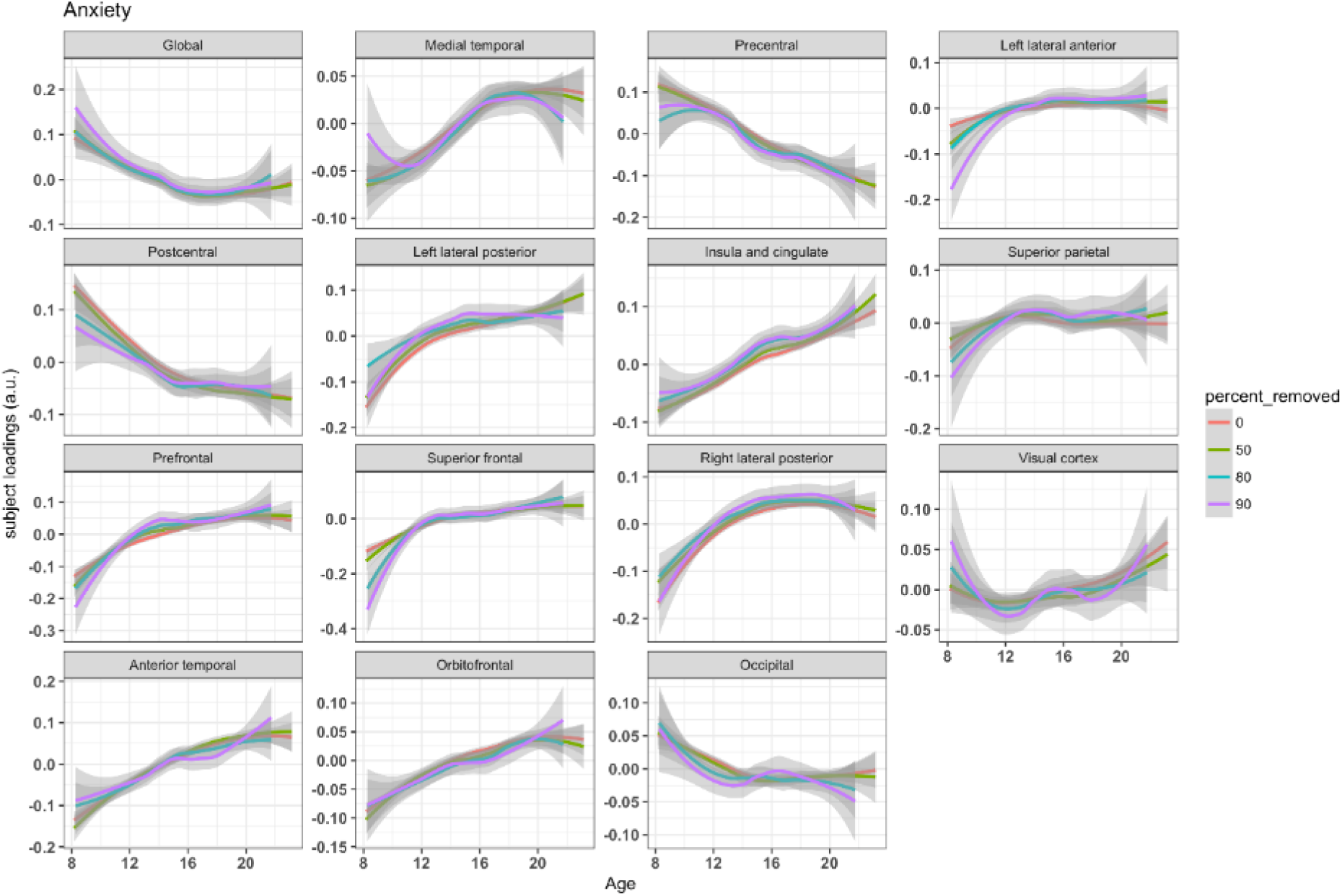
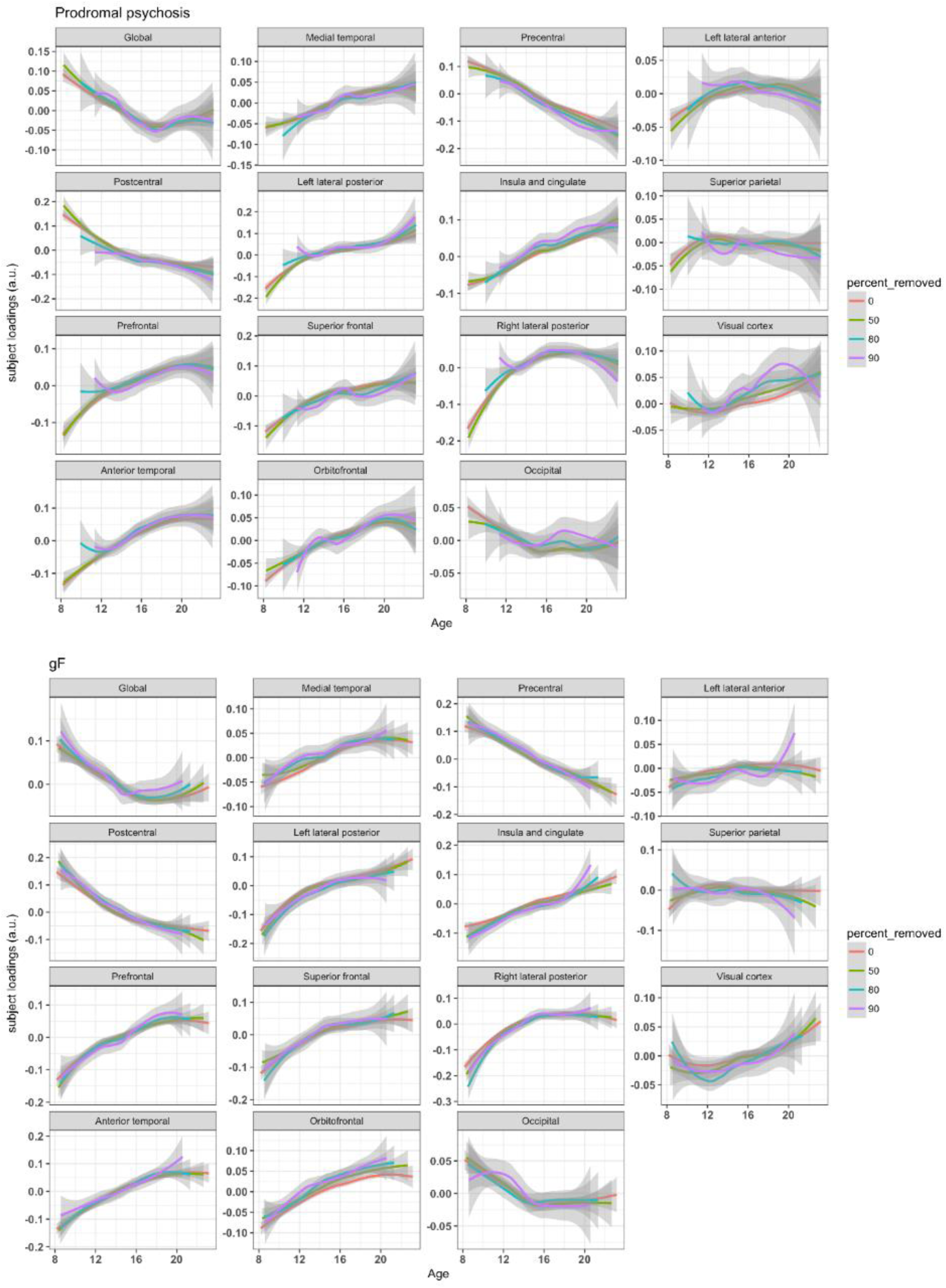
Loess subgroups visualization investigating the age-dynamic between psychopathology and GWC. The plots show the fitted curves for each GWC independent components (IC) for anxiety at the top, prodromal psychosis in the middle and gF at the bottom. The y-axis depicts the GWC IC loading, while the x-axis depicts age. The red line represents the full sample. The green, blue and purple line represents identical analyses after removal of individuals with the 50% 80% and 90% lowest psychopathology/gF loadings respectively. Shaded gray regions represent the confidence interval.

**Figure S7.**
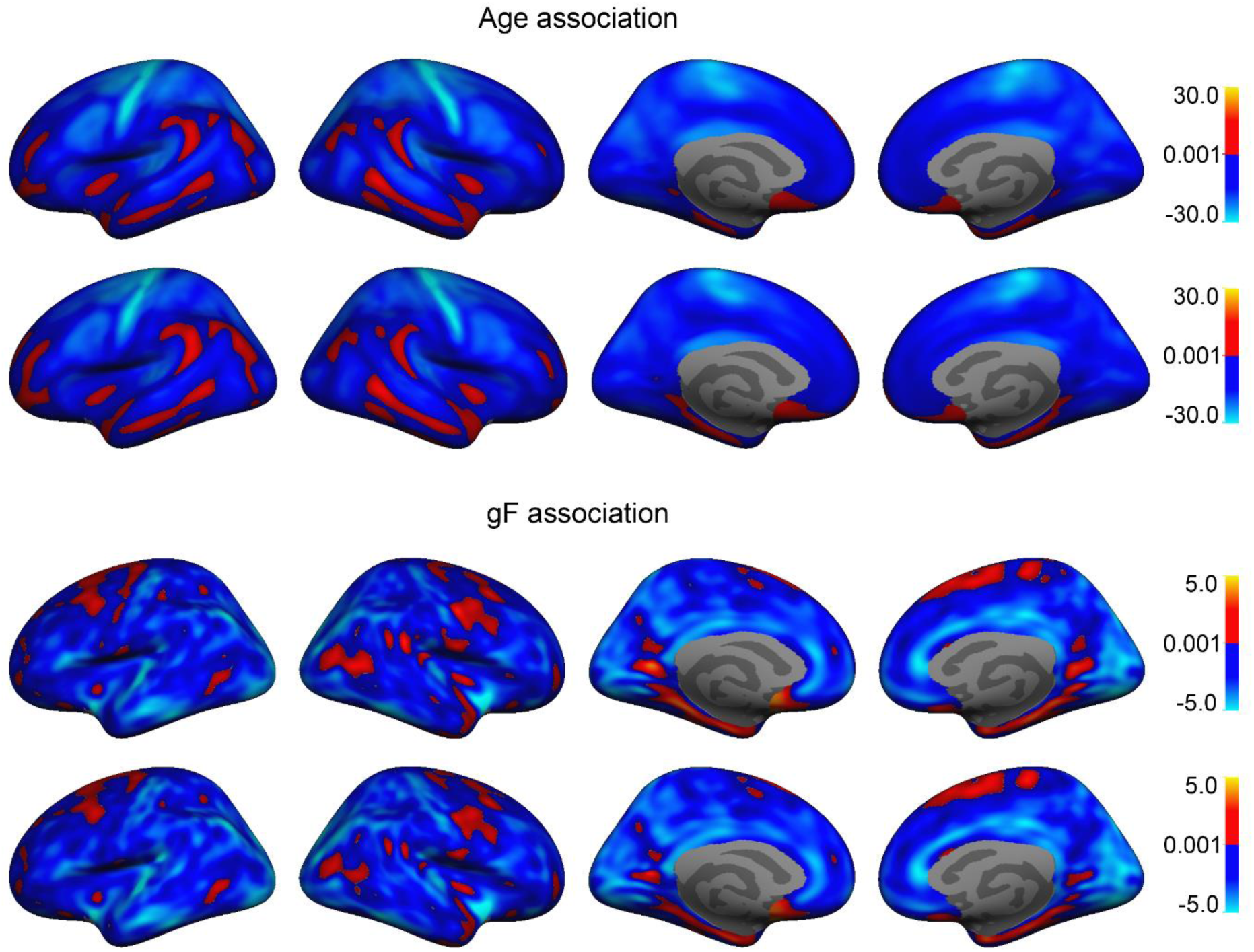
T statistic maps of vertex wise comparison of original results and results where thickness was added as a per-vertex co-variate. For the association between GWC and age (top two panels), the top panel shows our original t-stat surface map, while the bottom shows t statistics after adding thickness as a per-vertex co-variate. For the association between GWC and gF (bottom two panels), the top panel shows our original t-stat surface map, while the bottom shows t statistics after adding thickness as a per-vertex co-variate. Blue regions represent a negative age association, while red regions depict a positive association.

**Figure S8.**
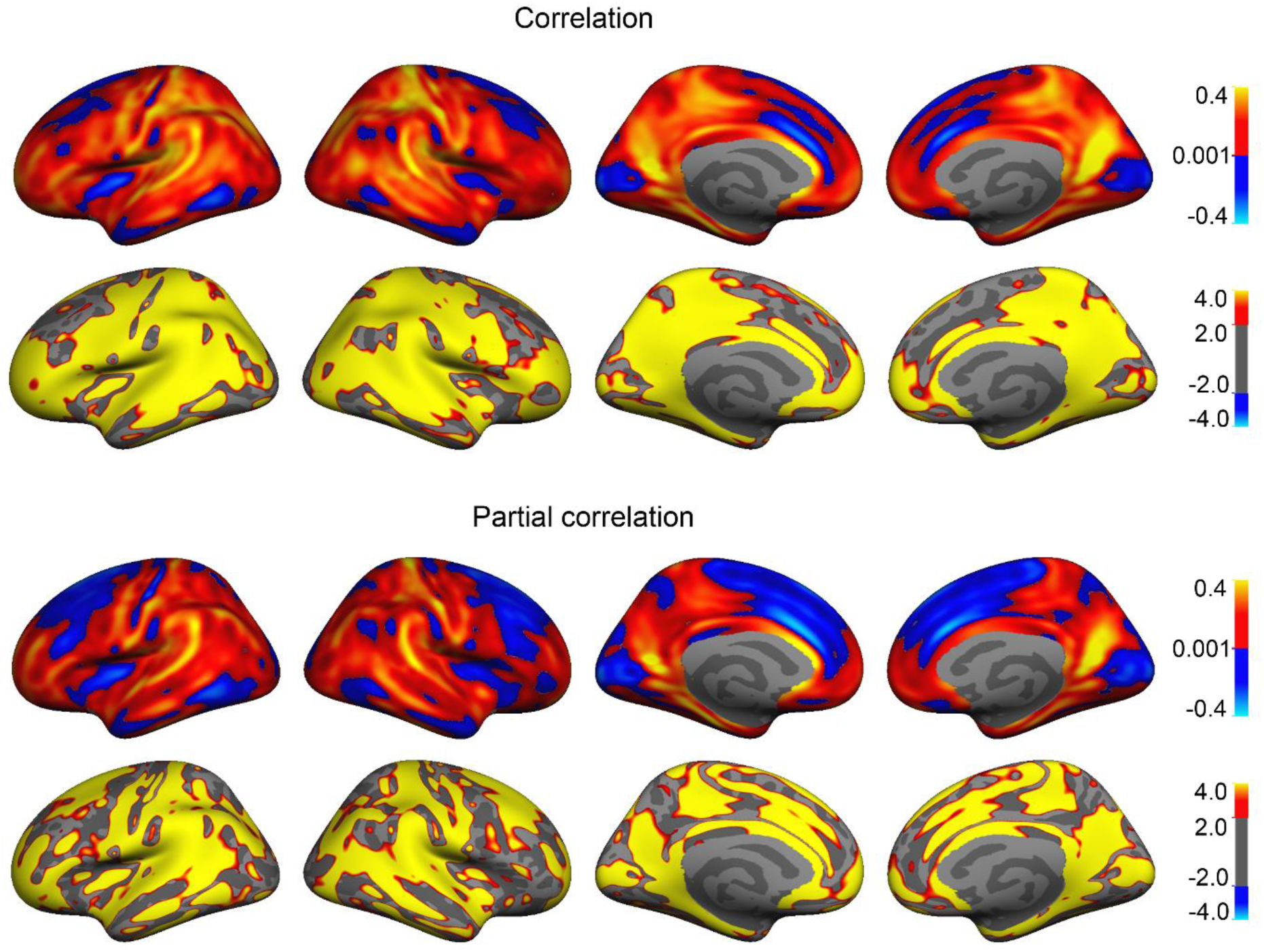
GWC and cortical thickness vertex wise full- and partial correlations after controlling for age. The top two panels show the vertex wise correlation, and corresponding FDR corrected p < 0.05 significance map converted to –log(p). The bottom two panels show the vertex wise partial correlation after controlling for age, and corresponding FDR corrected p < 0.05 significance map –log(p).

**Figure S9.**
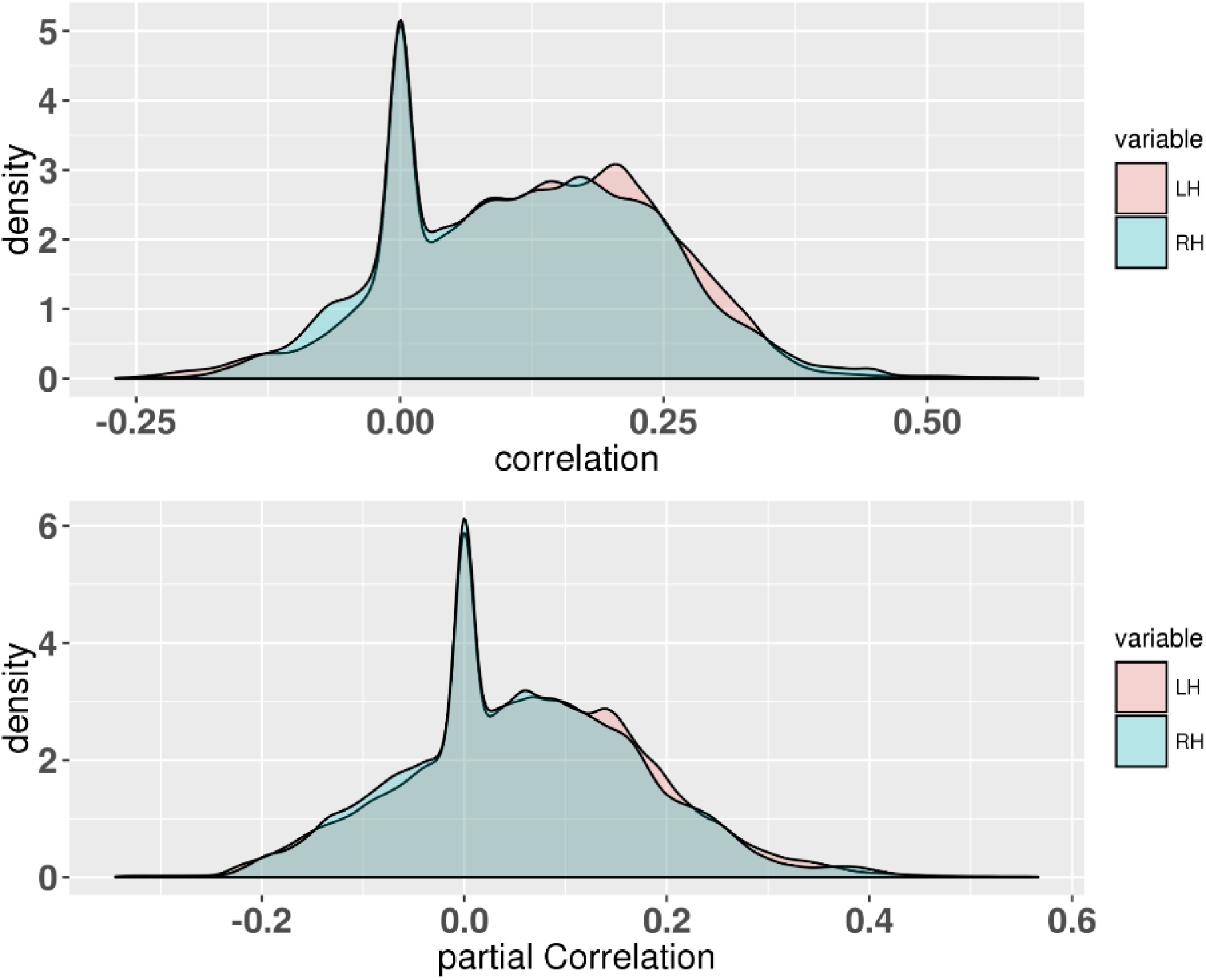
Density plot of GWC and cortical thickness vertex wise full- and partial correlations after controlling for age.

**Figure S10.**
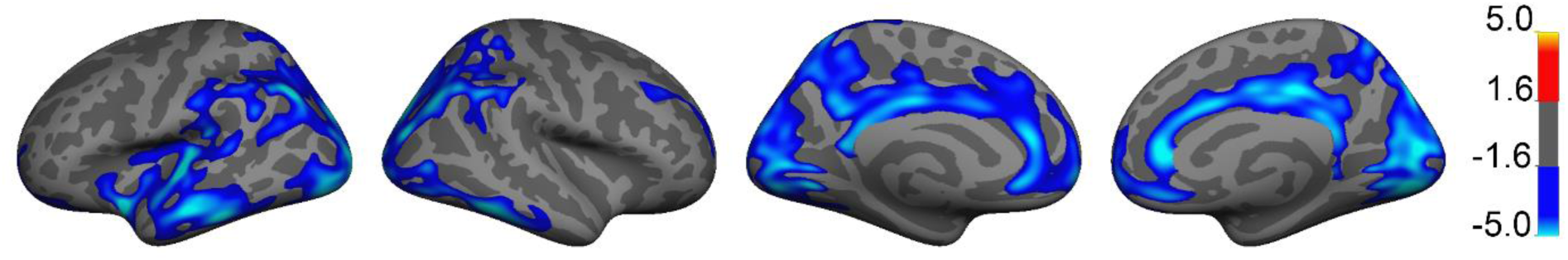
Vertex wise associations between gF and GWC. Permutation Analysis of Linear Models (PALM) on the associations of gF on GWC was performed. T statistics are masked by FWE corrected p-values thresholded at a minimum –log(p) of 1.6 to correct for two hemispheres. Blue regions represent a negative association, while red regions would depict a positive association.

**Figure S11.**
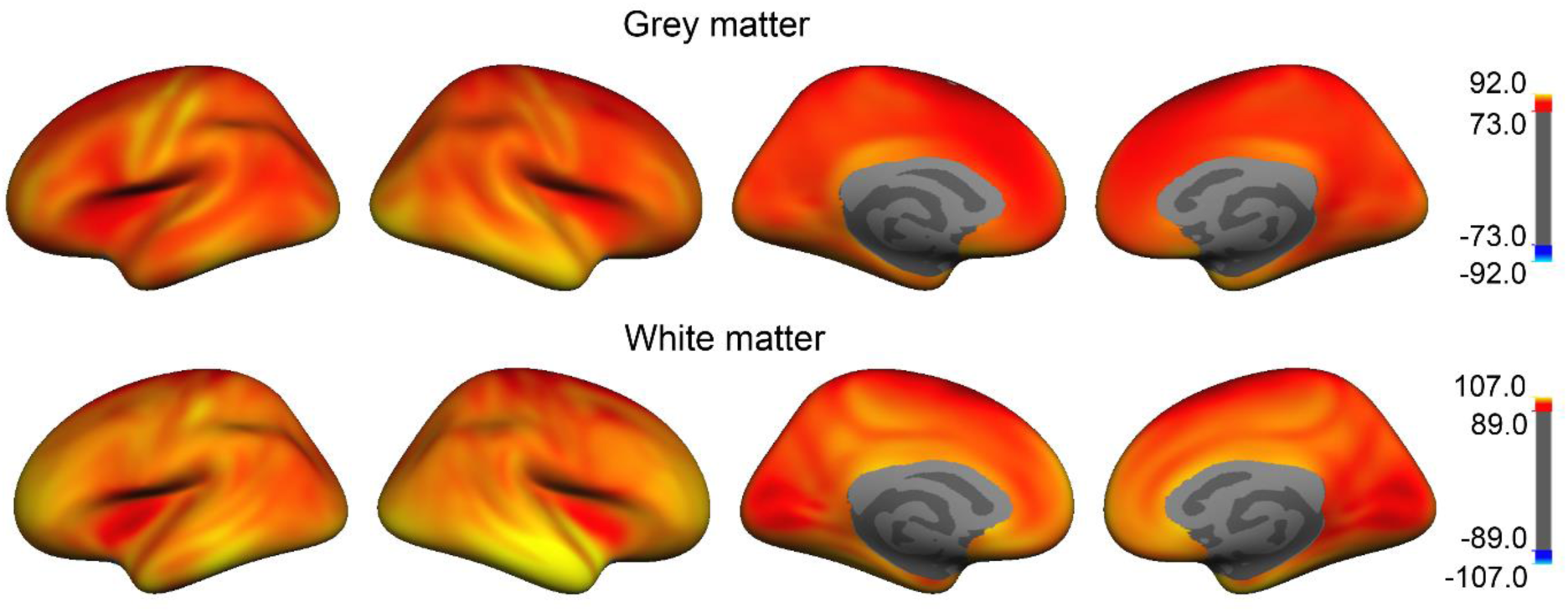
Gray matter intensity and white matter intensity mean surfaces. The top panel shows the mean concatenated gray matter surface of all subjects. The bottom panel shows the mean concatenated white matter surface of all subjects. The color bars show the intensities, and maps were smoothed using a Gaussian kernel of 10 mm full width at half maximum.

